# Tuning aromatic contributions by site-specific encoding of fluorinated phenylalanine residues in bacterial and mammalian cells

**DOI:** 10.1101/2022.04.12.488046

**Authors:** Grace D. Galles, Daniel T. Infield, Colin J. Clark, Marcus L. Hemshorn, Shivani Manikandan, Frederico Fazan, Ali Rasouli, Emad Tajkhorshid, Jason D. Galpin, Richard B. Cooley, Ryan A. Mehl, Christopher A. Ahern

**Affiliations:** Department of Molecular Physiology and Biophysics, University of Iowa College of Medicine, Iowa City, IA, United States; The GCE4All Research Center, Department of Biochemistry & Biophysics, Oregon State University, OR, United States; Theoretical and Computational Biophysics Group, NIH Center for Macromolecular Modeling and Bioinformatics, Beckman Institute for Advanced Science and Technology, University of Illinois at Urbana-Champaign

## Abstract

The aromatic side-chains of phenylalanine, tyrosine, and tryptophan interact with their environments via both hydrophobic and electrostatic interactions. Determining the extent to which these contribute to protein function and stability is not possible with conventional mutagenesis. Serial fluorination of a given aromatic is a validated method *in vitro* and *in silico* to specifically alter electrostatic characteristics, but this approach is restricted to a select few experimental systems. Here, we report a new group of pyrrolysine-based aminoacyl-tRNA synthetase/tRNA pairs that enable the site-specific encoding of a varied spectrum of fluorinated phenylalanine amino acids in *E. coli* and mammalian (HEK 293T) cells. By allowing the cross-kingdom expression of proteins bearing these unnatural amino acids at biochemical scale, these tools will enable deconstruction of biological mechanisms which utilize aromatic-pi interactions in structural and cellular contexts.

**Statement of Significance:** The aromatic side-chains of phenylalanine, tyrosine, and tryptophan are crucial for protein function and pharmacology due to their hydrophobic and electrostatic contributions to catalytic centers and ligand-binding pockets. However, few experimental approaches can chemically assess the functional roles of aromatics in cellular environments. The accepted computational method for aromatic interrogation is via serial fluorination, which lacks an experimental correlate in bacterial or mammalian cell systems. We have identified a family of synthetases to encode multiple different types of fluorinated phenylalanine residues in *E. coli* and HEK cells via nonsense suppression. The efficiency of these synthetases is sufficient to support biochemical characterization and structural determination of proteins with site-specific incorporation of unnatural phenylalanine analogs.

## Introduction

While primarily appreciated for its hydrophobicity, the benzyl side chain of phenylalanine also engages in various types of energetically favorable aromatic interactions (Figure 1A) [1-11]. Energetically significant aromatic interactions involving phenylalanine are believed to be widespread in nature [12, 13]. Based on emerging structural data, they have been increasingly proposed to play important mechanistic roles in ligand recognition [14-19], and protein-protein interactions, both in normal function [20, 21] and as a result of clinical mutation [22]. They may also play important roles in membrane anchoring, via attraction of the benzyl side chain to choline lipid headgroups [23], and have recently been proposed to mediate conduction in some ion channels [24]. Testing the specific functional role for aromaticity of the benzyl side chain in an interaction is not possible using conventional mutagenesis since mutation to a non-aromatic natural amino acid also causes concomitant changes in amino acid shape, convoluting data interpretation. Fluorination of the aromatic ring, on the other hand, allows for redistribution of the electrostatic potential of Phe with only minor steric changes (Figure 1B); thus allowing the specific testing of the functional relevance of aromaticity. Increasing the number of fluorine substitutions results in a linear decrease in the electrostatic potential for pi interactions [25, 26].

**Figure 1:**
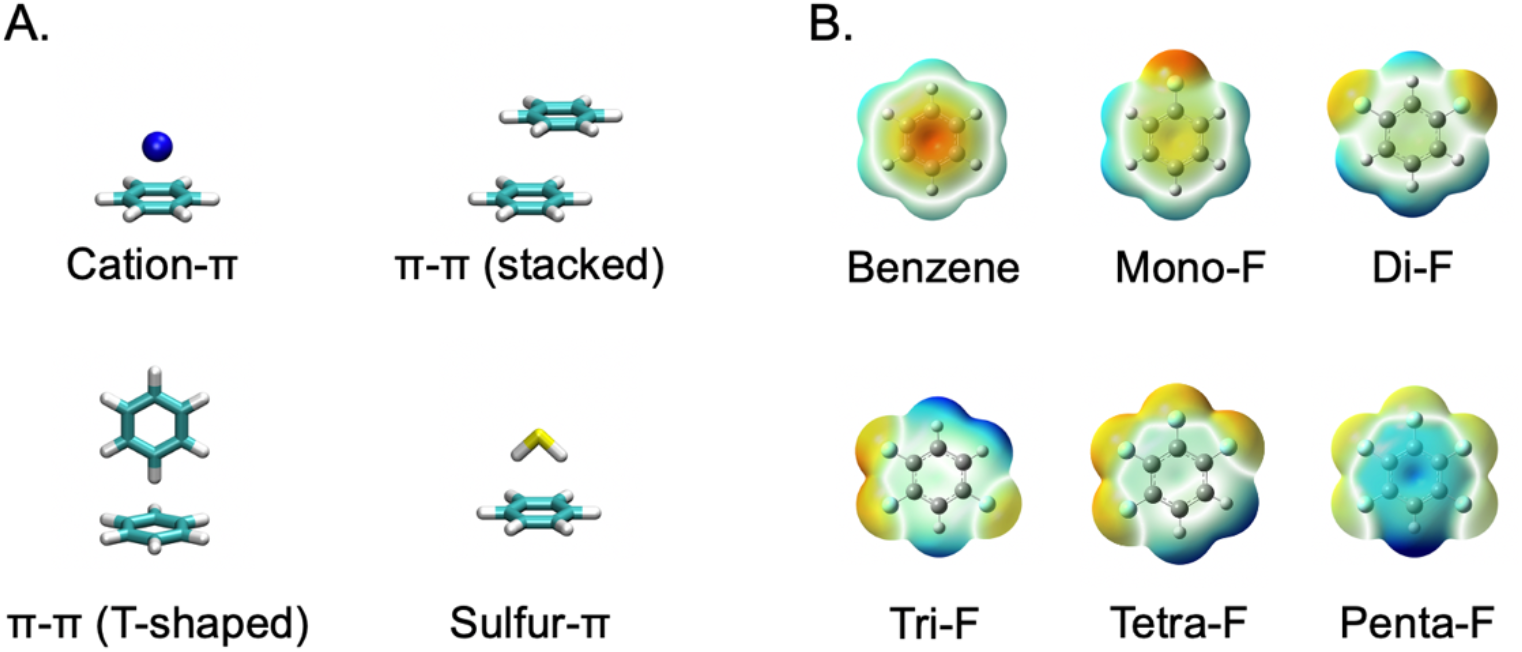
**A**. Illustrations of types of pi interactions in which Phenylalanine can participate. **B**. Electrostatic views of benzene (as a structural surrogate for Phenylalanine) and fluorinated analogs. Coloring is to convention of Red = negative, Blue = positive.

The site-specific installation of fluorinated phenylalanine amino acids can be accomplished via peptide synthesis and by the direct injection of misacylated nonsense suppressor tRNA [27], the latter of which has been used to functionally validate and quantify aromatic interactions in some membrane proteins, including ion channels [28-33]. These methods, while chemically flexible, cannot be easily scaled for biochemical and structural approaches. Additionally, some natural amino acids have been replaced with fluorinated, non-canonical structural analogs via permissive endogenous synthetases, but these methods are not site-specific, resulting in multiple sites within the protein sequence being modified heterogeneously [34]. On the other hand, nonsense suppression via coevolved tRNA / aminoacyl tRNA-synthetase (tRNA/RS) pairs allows for biochemical-scale production of proteins bearing site-specifically installed non-canonical amino acids (ncAA) in diverse cell systems and even organisms [27, 35]. Previously, one such tRNA/RS pair that encodes select fluorinated phenylalanine analogs into proteins was selected by rational design followed by screening of a single-residue library of 20 variants. This system was shown functional in *E. coli* if grown in minimal media without exogenous phenylalanine [36] but the lack of a sufficiently efficient tRNA-synthetase/tRNA pair stymies field progress. A method to accurately encode fluorinated Phe ncAAs in complex media and in eukaryotic cells would prove highly valuable for many clinically relevant soluble and membrane protein targets, including the myriad genes that must be expressed and/or studied in mammalian cells [12].

We hypothesized that by screening a greater number of tRNA-synthetase active site variants (~10^7^), using a novel strategy that enabled enrichment of variants able to selectively incorporate fluorinated phenylalanine derivatives but not phenylalanine, we may identify synthetases with the desired utility and versatility. We describe here the identification and validation of a group of synthetases, which we have named Phex, from the pyrrolysine system [37-39] which are competent for site-specific encoding of fluorinated di-, tri-, tetra, and penta-fluoro phenylalanine analogs within proteins in both bacterial and mammalian expression systems. Intact protein mass spectrometry defines the sythetase and ncAA combinations that enable high fidelity encoding at the amber suppression site without spurious substitution at natural codons. Finally we demonstrate biochemical scale encoding in two large (>150 KDa) membrane proteins of high clinical interest.

## Results

### A Pyl-based screen to identify synthetase variants that encode fluorinated phenylalanine analogs in E. coli

Historical attempts by our groups to screen for robust Pyl synthetases able to incorporate fluorinated phenylalanine analogs were not successful, possibly owing to the close steric resemblance of fluoro-benzene variants. In the present work, we decided to screen a library of Pyl synthetases with para-methyl, tetra-fluoro-phenylalanine (Figure 2A), in the hope that the additional structural bulk of the methyl at the para position would permit identification of synthetases selective against natural amino acids but selective for fluorinated Phe derivatives, perhaps including those lacking the para-methyl group. Two independent rounds (Figure 2B) of positive and negative screening within a Pyrrolysine-based aminoacyl tRNA synthetase library (see methods) in *E. coli* [40] resulted in the identification of several synthetases that enabled amber suppression within superfolder

**Figure 2:**
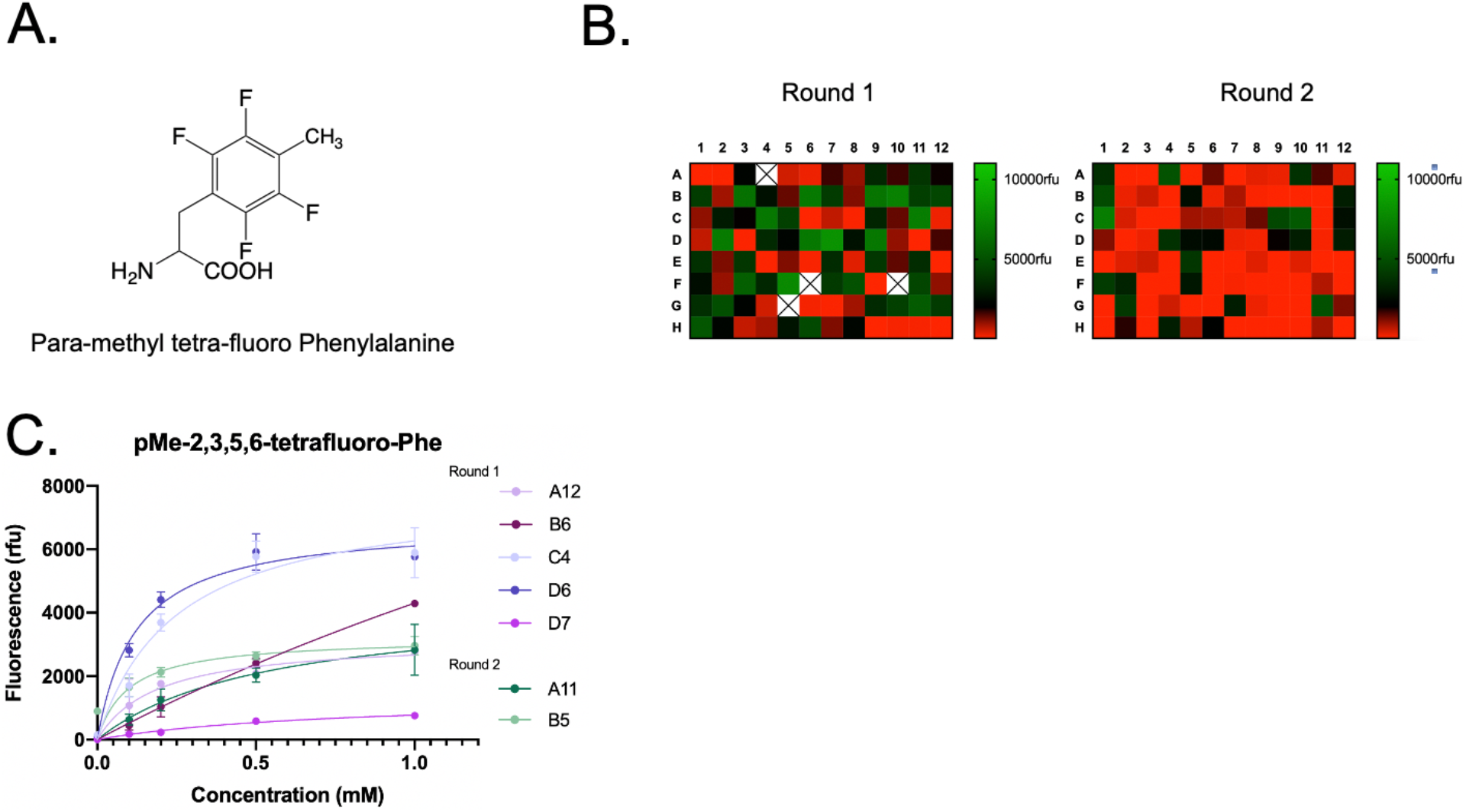
Strategy and identification of synthetases. **A**. Chemical structure of para-methyl tetra-fluoro Phe, which was used for screening in a PylRs library in *E. coli*. **B**. Heatmap of synthetases hits that resulted from positive and negative screening in terms of their relative GFP N150TAG rescue (fluorescence) in the presence of para-methyl tetrafluoro Phenylalanine. **C**. Unnatural Protein 50 (UP50) curves for the top performing synthetases from the screens.

GFP (sfGFP_N150TAG) in response to para-methyl tetra-fluoro Phe (Figure 2B). Unique synthetases wherein suppression of sfGFP (from GFP signal) was greater than 10-fold above background are shown in Figure 2C. As a preliminary assessment of the quality of the evolved tRNA/aminoacyl synthetase (RS) pairs, experiments to determine the concentration of ncAA required to express ncAA-sfGFP at half its maximum expression level, called the Unnatural Protein 50% (UP50), [40] were performed for the synthetases using para-methyl tetra-fluoro Phe (Figure 2C). The UP50 values for para-methyl tetra-fluoro Phe for the selected tRNA/RS pairs validate they will function efficiently at 0.5 mM ncAA [40].

### Synthetases selected for para-methyl-tetra-fluoro-Phe are permissive to a panel of

*fluorinated Phe analogs*

An important purpose of the screening efforts was to identify synthetases that enable the encoding of fluorinated (but non-methylated) phenylalanine analogs within proteins. Therefore we tested for permissivity [41] of the selected synthetases against a diverse collection of phenylalanine analogs (Figure 3A). In practice, this was an iterative process, with preliminary results from both *E*.*coli* (Figure 3) and HEK cells (discussed below) motivating the synthesis and evaluation of additional structural analogs to audition in both systems. Interestingly, the synthetases tested encoded penta-fluoro Phe and 2,3,5,6 tetra-fluoro Phe at comparable levels to para-methyl tetra-fluoro Phe in *E. coli*, indicating that the para-methyl group is not required for recognition (Figure 3B). Each synthetase showed some degree of permissivity; Phe_X-B5_ and Phe_X-D6_ were particularly interesting examples with substantially different recognition profiles (Figure 3B). Phe_X-B5_ displayed broader utility, but also lower apparent specificity over background, than Phe_X-D6_ in *E. coli*.

**Figure 3:**
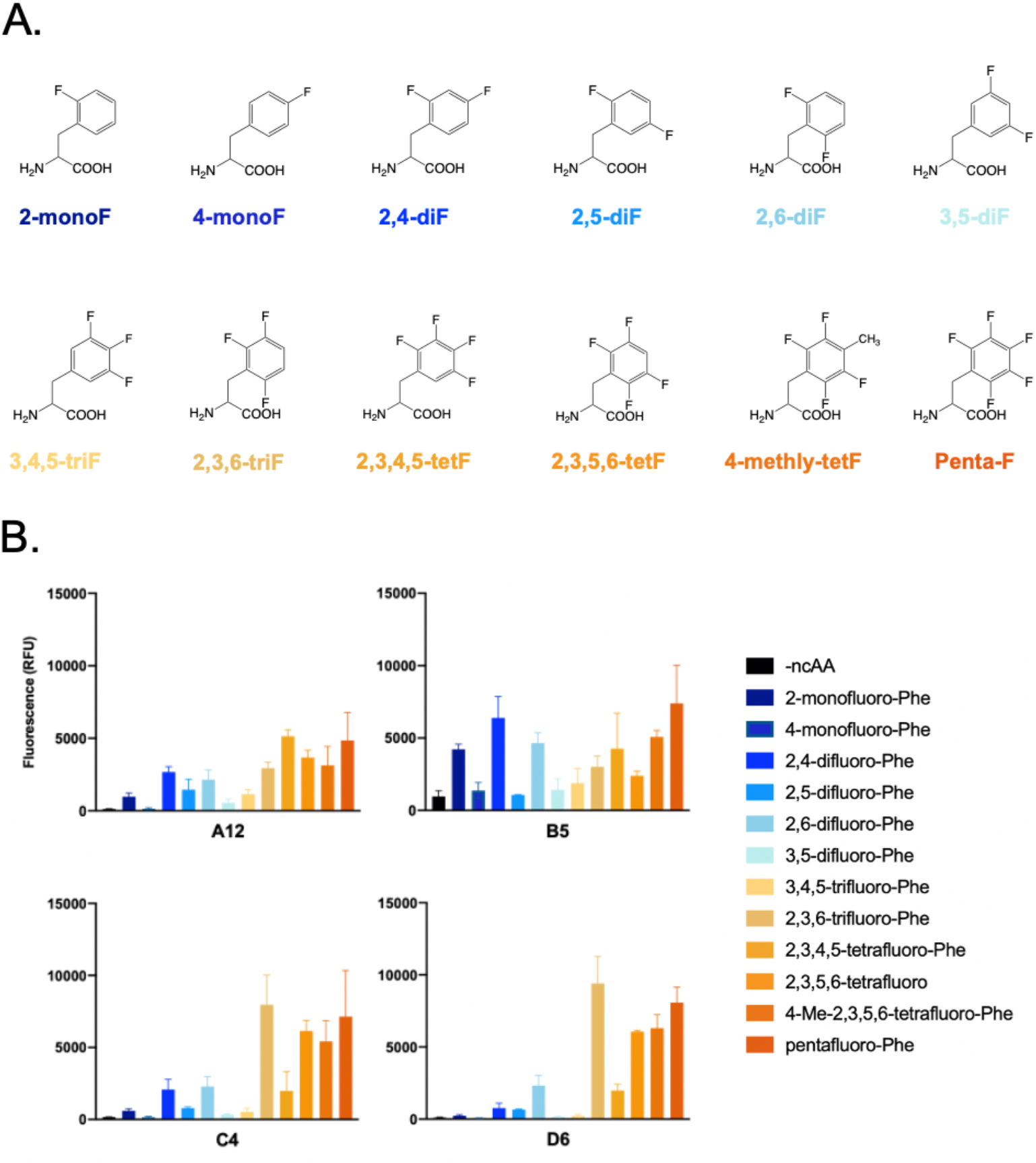
Permissivity of the identified synthetases. **A**. Chemical structures of the fluorinated phenylalanine residues used in the screen. **B**. Green fluorescence from rescue of sfGFP-N150TAG in *E*.*coli* using 1 mM of each ncAA after 24-hour incubation.

With permissivity profiles in hand, we performed ab-initio quantum calculations to determine the effect of specific fluorination patterns on the potential of each analog to encage in electrostatic interactions, which has been previously done for a few of these analogs [25]. We calculated the theoretical ΔG (kcal/mol) for interactions with three different cations and their representative benzene structures (Table 1). As expected, increasing the number of fluorine atoms elicited a linear decrease in the binding energy. Table 1 expresses both the derived ΔG values and how they compare as a percentage of the native benzene of Phe (% Phe). Importantly, the relative positioning of the fluorine atoms did not greatly affect energetics of binding, suggesting that the synthetases encode analogs with different potential local electrostatic effects but equivalent global perturbations. The encoding of multiple di-, tri-or tetra-fluoro Phe analogs may prove useful to better understand phenylalanine side chains involved in complex, or multivalent motifs [25, 42].

**TABLE 1:** Effects of fluorination on interaction energies with cations

|  |  |  | Mono- |  | Di- |  |  |  | Tri- |  |  |  | Tetra- |  |  |  | Penta- |  |
| --- | --- | --- | --- | --- | --- | --- | --- | --- | --- | --- | --- | --- | --- | --- | --- | --- | --- | --- |
| Benzene analog | Benzene |  | 1F |  | 1,3F |  | 1,4F |  | 1,2,3F |  | 1,2,5F |  | 1,2,3,4F |  | 1,2,4,5F |  | 1,2,3,4,5F |  |
| analogous Phe<sub>x</sub> substrate(s) | Phe |  | 2F |  | 2,4F, 2,6F |  | 2,5F |  | 3,4,5F |  | 2,3,6F |  | 2,3,4,5F |  | 2,3,5,6F |  | 2,3,4,5,6-F |  |
|  | ΔG (kcal/mol) | % PHE | ΔG (kcal/mol) | % PHE | ΔG (kcal/mol) | % PHE | ΔG (kcal/mol) | % PHE | ΔG (kcal/mol) | % PHE | ΔG (kcal/mol) | % PHE | ΔG (kcal/mol) | % PHE | ΔG (kcal/mol) | % PHE | ΔG (kcal/mol) | % PHE |
| Sodium | 26.7 | 100 | 22.8 | 85.4 | 19.1 | 71.5 | 18.7 | 70.0 | 16.2 | 60.7 | 15.6 | 58.4 | 12.7 | 47.6 | 11.9 | 44.6 | 9.1 | 34.1 |
| Ammonium | 19.5 | 100 | 16.3 | 83.6 | 13.1 | 67.2 | 13 | 66.7 | 10.2 | 52.3 | 10.1 | 51.8 | 7.4 | 37.9 | 6.8 | 34.9 | 4.6 | 23.6 |
| Guanidinium (T-shaped) | 14.8 | 100 | 12.3 | 83.1 | 10.3 | 69.6 | 9.7 | 65.5 | 8.1 | 54.7 | 6.2* | 41.9 | 3.9* | 26.4 | 3.9* | 26.4 | 1.8* | 12.2 |

\*derived from single point calculation

We next verified the fidelity of the top tRNA/RS pairs by analyzing the intact sfGFP proteins by ESI-mass spectrometry to confirm accurate encoding of the fluorinated phenylalanine derivatives [40]. We used the Phe_X-D6_ and Phe_X-B5_ synthetases to express and purify sfGFP_N150TAG_HIS in *E. coli*. Whole protein sfGFP masses were consistent with replacement of N150 with each of the respective F-Phe ncAAs. Using Phe_X-D6_, we observed evidence of high-fidelity incorporation (Figure 4) for di-, tri-, tetra- and pentafluoro phenylalanine (*see* methods for additional details). When the Phe_X-B5_ synthetase was used to express sfGFP bearing a range of ncAAs at N150, evidence of efficient encoding was clear for a wide range of fluorinated ncAAs (Figure 4). However, for several ncAAs with lower encoding efficiency, detectable peaks corresponding to the mass of Phe at N150 were observed (Fig 4, Δ mass of +33 Da from asparagine, asterisks). This background encoding of natural AAs was expected from the higher fluorescence values in the absence of ncAA for Phe_X-B5_ (Figure 3B) suggesting that Phe_X-B5_ incorporates natural amino acids in *E. coli* when using ncAAs poorly recognized by the synthetase.

**Figure 4:**
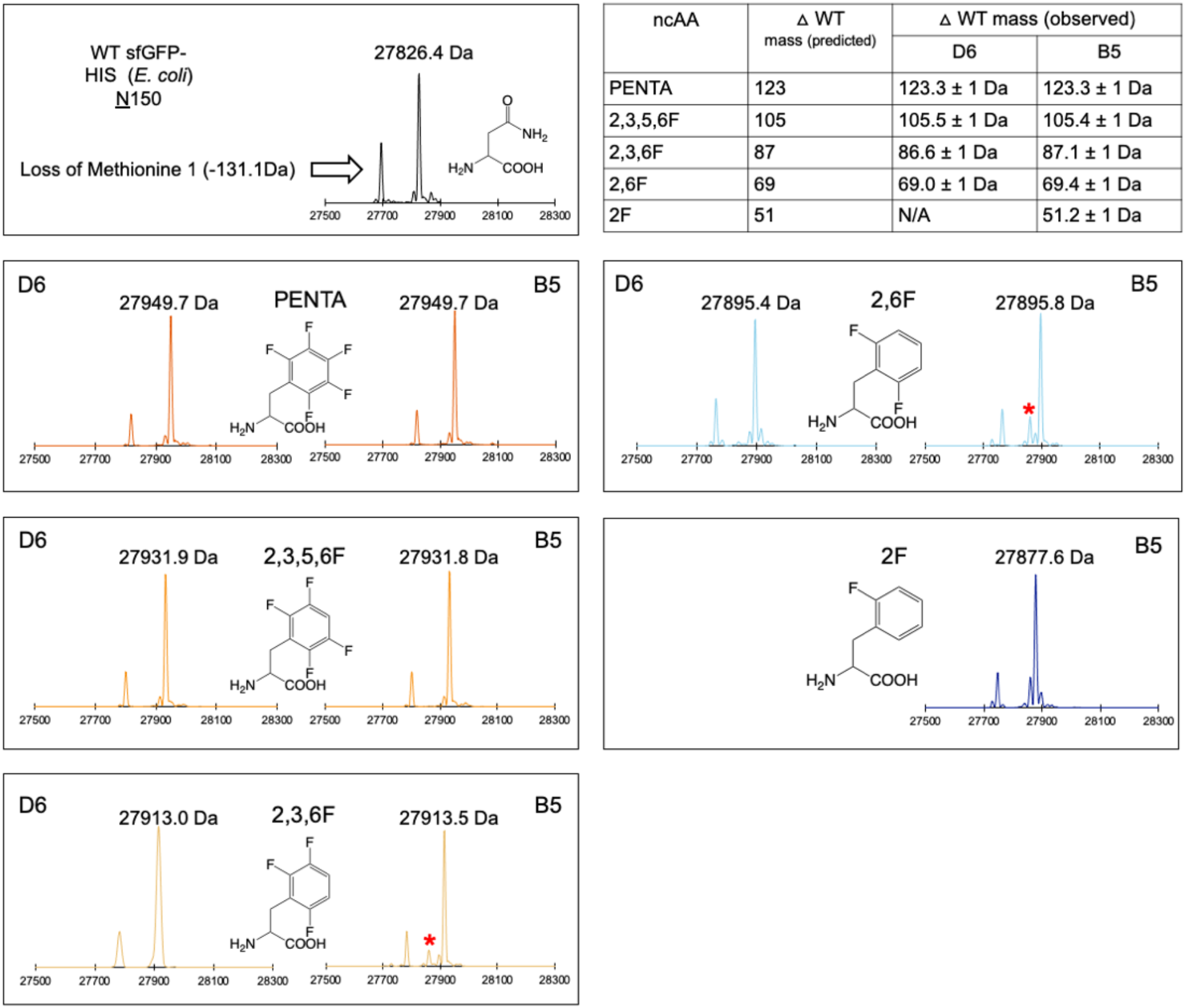
ESI mass spectrometry confirmation of sitespecific encoding of fluorinated Phe residues within superfolder GFP in *E coli*. sfGFP was purified via His tag from 50 mL growths. Table at top right: mass differences of sfGFP N150TAG from WT GFP conform to prediction for the molecular mass of each fluoro Phe species. In some conditions for B5 synthetase, a small fraction of mass consistent with Phe incorporation was observed (asterisks). Mass spectrometry was not pursued for the combination of 2F Phe and D6 due to low expression in the screen.

### Site-specific encoding of fluoro-Phe ncAAs into proteins in HEKT cells

To evaluate the encoding of fluorinated analogs in HEK cells, we cloned the Phe_X-B5_ and Phe_X-D6_ synthetases into the pAcBac1 [43] plasmid and co-transfected them with a plasmid encoding sfGFP_N150TAG in the presence or absence of fluoro-Phe ncAAs. In HEK cells both the Phe_X-D6_ and Phe_X-B5_ tRNA/RS pairs produced sfGFP in the presence of fluoro-Phe ncAAs but not appreciably in their absence, yielding green cells which were imaged for fluorescence at 24 hours (Figure 5A). For quantitative comparison of ncAA-protein expression, fluorescence from cell lysate was measured. As with *E. coli* fluoro-Phe encoding, the position and number of fluorine substitutions affected the efficiency of producing ncAA-protein with the Phe_X-D6_ and Phe_X-B5_ tRNA/RS pairs (Figure 5B). Of note, the relative fidelity of these tRNA/RS pairs was better in HEK cells compared to that in *E. coli*., but the background fluorescence from the Phe_X-B5_ tRNA/RS pair was still higher than Phe_X-D6_ (estimated 0.84% and 0.15% of maximum encoding for Penta-F Phe for Phe_X-B5_ and Phe_X-D6_ pairs, respectively).

**Figure 5:**
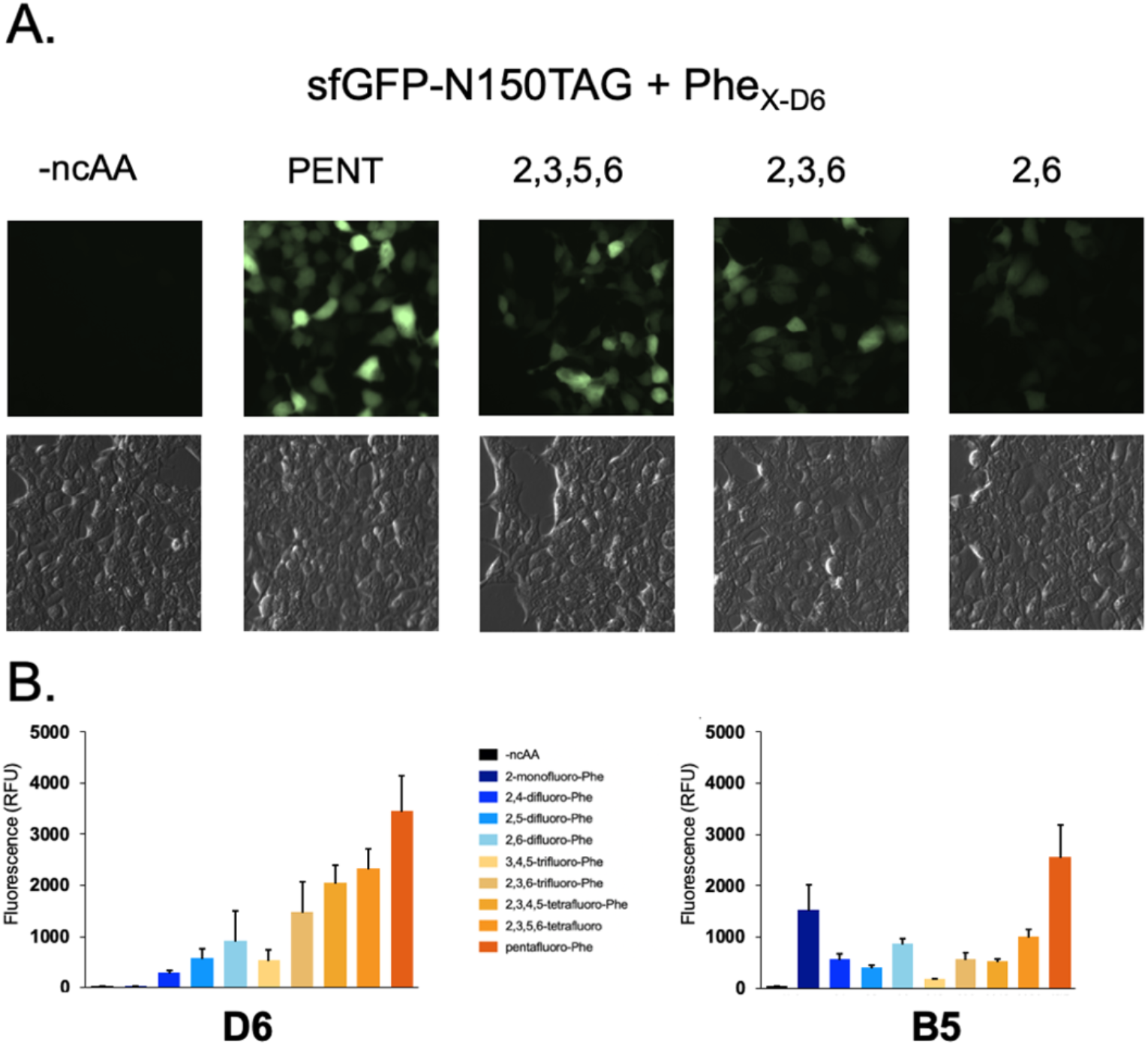
Function of Phex synthetases in HEKT cells. **A**. Micrographs of HEKT cells co-transfected with GFP_N150TAG and Phe_x.D6_in the presence or absence of the given amino acid. GFP (top) and brightfield (bottom). In all cases, images taken approximately 24 hours after transfection and unnatural amino acid used at 2 mM concentration. **B**. Lysate GFP fluorescence for a range of fluorinated phenylalanine residues.

To rigorously confirm the efficiency and fidelity of these tRNA/RS pairs in a eukaryotic context, ncAA-sfGFP was expressed in HEKT cells at 2 mM ncAA, purified via c-terminal His_6_ tag and subjected to ESI-mass spectroscopy analysis. The Phe_X-D6_ and Phe_X-B5_ pairs encoded multiply fluorinated Phe analogs with high fidelity (Figure 6). Estimated yield of sfGFP with Penta-fluoro Phe encoded by Phe_X-D6_ was ~34 μg per gram of HEK cells (Supplemental Fig 1). From the *E. coli* permissivity evaluation we note a primary feature distinguishing Phe_X-B5_ from Phe_X-D6_ is the ability of the former to recognize 2-mono fluoro Phenylalanine. However, in HEKT cell expression of sfGFP-N150TAG with Phe_X-B5_ and 2-mono fluoro Phe, we observed evidence of widespread encoding of 2-mono fluoro Phe at Phe-codons (Supplemental Fig 2). This observation suggests that the endogenous (human) Phe tRNA/RS pair cannot effectively distinguish between 2-mono F Phe and Phe, preventing site-specific encoding of 2-monofluoro Phe in HEKT cells and likely other human cells as well. Overall, the ESI-mass spectroscopy analysis of the pure ncAA-protein demonstrate the fidelity of the Phe_X_ RS group to encode 6 of the fluorinated ncAAs; 2 di-fluorinated (2,5/2,6), a tri-fluorinated (2,3,6), two tetra-flourinated (2,3,5,6/2,3,4,5), and the penta-fluorinated phenylalanine in HEKT cells with high efficiency and fidelity. Dose-response experiments with ascending concentrations of the ncAAs indicate that a few combinations (such as Phe_X-D6_ + penta-fluoro Phe) are useable at sub-millimolar concentrations of ncAA but that, in general, 1-2 mM is necessary for efficient ncAA-protein expression (Supplemental Fig 3).

**Figure 6:**
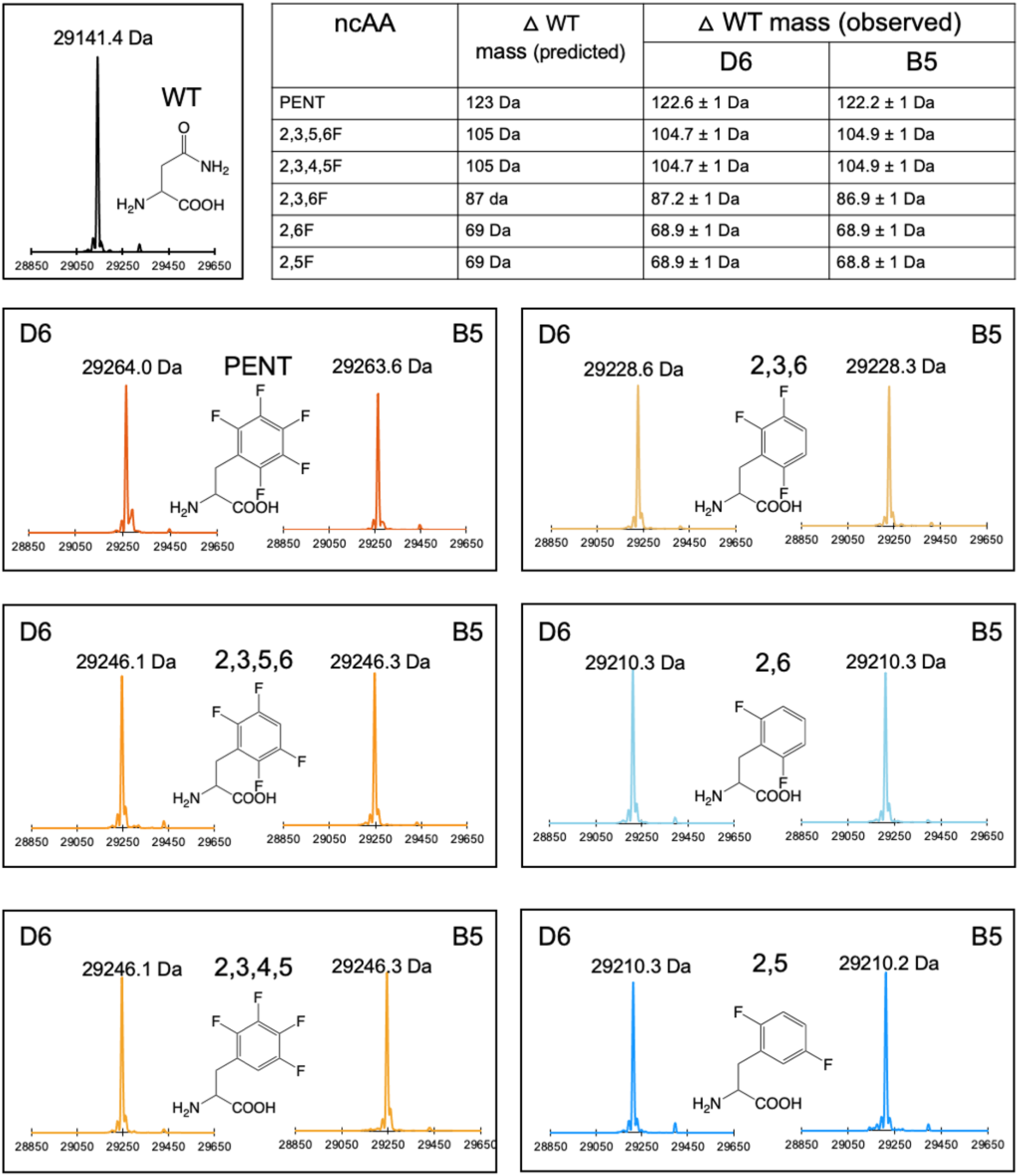
ESI mass spec confirmation of encoding of fluorinated Phe analogs within sfGFP expressed in HEKT cells. ncAAs were added at 2 mM concentration and cells were harvested approximately 36 hours post transfection. sfGFP-WT-V5-HIS or N150_TAG-V5-HIS was purified via HIS tag. Table (at top right) reports mass differences of sfGFP N150TAG from WT GFP, which conform to prediction for the molecular mass of each fluoro Phe species.

### Use of Phe_x_ synthetases to site-specifically encode fluoro-Phe ncAAs into human proteins

Finally, we demonstrate the utility of the tRNA/Phe_x_ for fluoro-Phe ncAA encoding within two large, human ion channels of high clinical significance. Mutations within the ~150 kDa Cystic Fibrosis Transmembrane Conductance Regulator (hCFTR) cause the life-shortening disease Cystic Fibrosis (CF) (Figure 7A) [44], while mutation and dysregulation of the ~200 kDa cardiac sodium channel (hNa_V_ 1.5) are associated with various cardiovascular diseases including cardiac arrythmia and heart failure (Figure 7B) [45, 46]. These proteins have been studied in a wide variety of eukaryotic cell hosts, but mammalian cell expression is either essential or preferred for many types of functional, biochemical, and structural analyses of channel regulation or pharmacology [47-60]. We introduced TAG codons at position F508 in CFTR and at F1486 in hNav 1.5 and co-transfected these expression plasmids with Phe_X-D6_ in the presence or absence of 2,3,6F Phe. We harvested the cells two days post transfection and assessed expression via western blot of cell lysates. In both cases, expression of full-length channels (as indicated by antibodies specific for C-terminal epitopes) was dependent on addition of 2,3,6F Phe in the growth media (Figure 7 A-C), demonstrating specific encoding by Phe_X-D6_. Independent confirmation of 2,3,6F Phe encoding within Na_v_ 1.5 was accomplished via expression of Na_v_ 1.5 F1486(2,3,6F Phe) via Phe_X-D6_, purification via FLAG TAG and in-gel tryptic digestion / MS-MS determination (Figure 7D). Finally, WT and Nav F1486(2,3,6F Phe) channels were expressed in HEK cells and recorded via the whole cell patch clamp configuration. In agreement with the western blot results, robust macroscopic currents were observed for both WT and mutant channels; these channels appeared to activate and inactivate normally in response to membrane depolarization (Figure 7E). Analysis of gating parameters revealed that encoding 2,3,6F Phe at position F1486 caused a subtle enhancement of channel inactivation. This was apparent in a small (~7 mV) but statistically significant left shift in the midpoint of steady state inactivation (Figure 7G, Supplemental Table 1). Interestingly, data from exhaustive natural mutation of this critical phenylalanine previously revealed a strong correlation between hydrophobicity and efficiency of inactivation; no natural mutation fully reproduced WT (Phe) behavior [61]. The data here suggest that the interaction of the IFM particle for its inactivated-state receptor [53] is enhanced via near-isosteric fluorine substitution on the benzene ring of F1486. This is consistent with studies of model hydrophobic cores wherein targeted fluorination enhanced thermodynamic stability over native levels [42].

**Figure 7.**
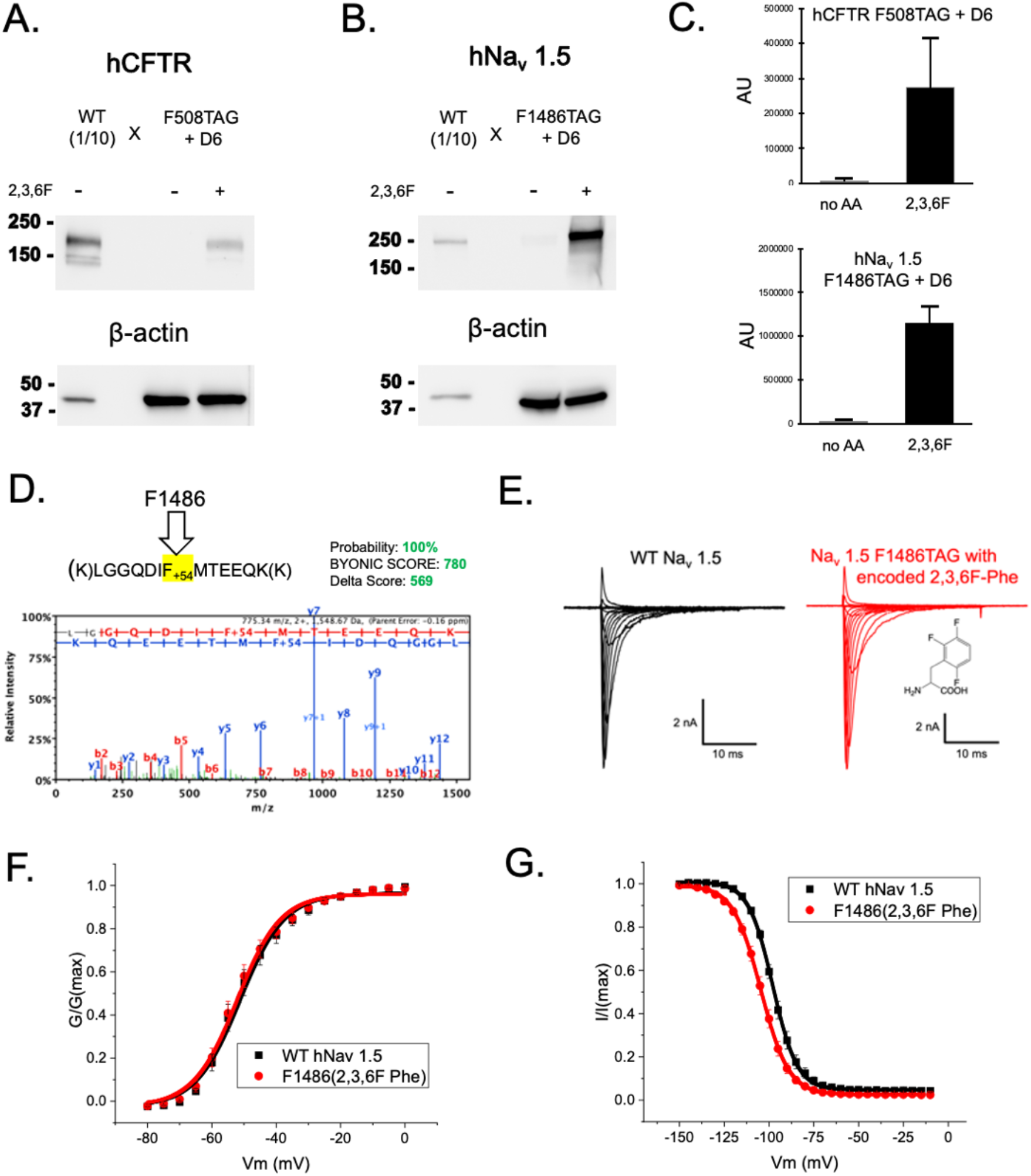
Site-specific encoding of tri-fluoro-Phe within two large, human ion channels. **A**. Western blot of CFTR using C terminal antibody (ab596) of WT CFTR or F508TAG-CFTR cotransfected with Phe_X-D6_ in the presence or absence of 2 mM 2,3,6 F Phe. WT lysates are loaded at 1:10 of TAG conditions **B**. Western blot of Nav 1.5 using C terminal antibody (D9J7S) of WT hNav 1.5 or F1486TAG-hNa_v_ 1.5 co-transfected with Phe_x.D6_ in the presence or absence of 2 mM 2,3,6 F Phe. **C**. Quantification (densitometry) of signal from 60 jig of loaded lysate. **D**. Mass spectra of tryptic fragment from expressed and purified hNav 1.5 F1486(2,3,6F Phe) via D6. E. Example current families for whole (HEK) cells expressing WT or mutant channels. Cells held at -140 mV and activated via depolarization in 5 mV increments. **F**. Activation (GA/) curves for WT and mutant channels. **G**. Steady state inactivation curves for WT and mutant channels.

## Discussion

The electrostatic contributions of aromatic residues to protein-protein interactions and ligand recognition are becoming increasingly appreciated [12], but available tools limit the ability to test proposed mechanistic hypotheses. The tRNA/ Phe_X_-RS pairs reported herein allow for the encoding of a wide spectrum of fluorinated phenylalanine residues, which can be used to tune the electrostatic contributions of the aromatic side chain in pi-pi and cation-pi interactions (Figure 1, Table 1). The majority of the fluorinated phenylalanine residues used in this study are commercially available, making this tool set immediately adoptable for those studying aromatic side change contributions in prokaryotes and eukaryotes. As these tRNA/RS pairs were first evolved for a multiply fluorinated para-methyl-Phe analog, it is not surprising that the more highly fluorinated species tend to encode most efficiently into proteins (Figure 3,5). We view the fact that penta-fluoro and tetra-fluoro phenylalanine analogs encode so well as fortuitous because it enables the investigator to gain valuable early information on the electrostatic role of Phe residues given the strong electrostatic perturbation imparted by the fluorines. For example, penta-fluoro phenylalanine is estimated to bear only ~12-34% of the cation-pi binding potential of native Phe (Table 1), whereas tri-fluorinated phenylalanine bears ~40-60%, depending on the cation involved.

The efficiency of the synthetases as demonstrated herein suggest that they will be useful for the site-specific encoding of non-canonical phenylalanine analogs within proteins in *E. coli* and HEK cells at levels sufficient for biochemical characterization. In HEK cells, the yield of the most robust combination (Phe_X-D6_ and penta-fluoro phenylalanine) was 34 µg per gram of cell pellet using transient transfection. This is comfortably within the range necessary for structure determination by cryo-EM, provided the protein is efficiently purified and the growth is sufficiently scaled. The system may be further optimized via construction of synthetase-incorporated baculoviruses [43] or stable cell lines. The function of the synthetases in mammalian cells enables their use to answer questions that were previously intractable with other systems (prokaryotic, cell-free, *Xenopus* oocyte) and in principle can be extended to express and characterize myriad membrane and soluble proteins. Beyond mechanistic work, the synthetases may be used to site-specifically incorporate fluoro-Phenylalanine NMR probes [62], and an optimized system utilizing these synthetases may enable the biotechnological production of therapeutic proteins wherein fluorinated phenylalanine residues are encoded for pharmacologically advantageous reasons [63].

## Methods

### Sources of Amino Acids

#### Synthesis of 2,3,5,6 tetrafluoro-paramethyl-Phe

Synthesis was carried out according to the report of Mikaye-Stoner et al. [64] Mass spectrometry revealed mass of 252.05 in positive mode and 250.06 in negative mode, which agreed with the predicted mass of 251.06.

#### Synthesis of 2,3,5,6 tetrafluoro Phe

All solvents and reagents were supplied by Sigma Aldrich and were used as-is unless explicitly stated. Dry nitrogen was supplied by Praxair and passed through two moisture scrubbing columns of dry calcium sulfate (Drierite) prior to use. HPLC analyses were performed on a Waters 1525 Binary HPLC pump equipped with a Waters 2998 Photodiode Array Detector, employing Sunfire C18 analytical (3.5 µm, 4.6 mm x 150 mm, 0.8 ml/min) or preparative (5.0 µm, 19 mm x 150 mm, 10 ml/min) columns and Empower software. Buffers were drawn in linear gradients from 100% A (50 mM ammonium acetate) to 100% B (acetonitrile) over 30 min. Mass spectra were recorded on a Waters QToF Premier Quadrupole instrument, in both positive and negative modes. The amino acid was synthesized according to a published procedure [42] with the following small modification: in lieu of ethanol extraction, following saponification of the methyl amide, the neutralized and lyophilized salt was purified directly via HPLC. The product was lyophilized to a white powder. (Calculated mass for C_9_H_7_F_4_NO_2_: 237.1 Da, found 238.0 Da (M+1)/236.0 Da (M-1) for MS in positive and negative modes).

The other Phe analogs were acquired commercially as follows**: 2F** Phe -Astatech, cat # 73308; **4F** Phe-Chem-impex cat # 02572; **2**,**4F** Phe-1ClickChemistry, cat #5C96757; **2**,**5F** Phe-Astatech, cat # 60350; **2**,**6F**-Phe HCl salt-Chem-impex, cat #24171; **3**,**5F** Phe Chem-impex, cat # 04123; **2**,**3**,**6F**-Phe-Astatech, A50355; **3**,**4**,**5F** Phe-Chem-impex, cat # 07394; **2**,**3**,**4**,**5F** Phe-Enamine, cat # en300-27751009; **2**,**3**,**4**,**5**,**6F** Phe-Chem-impex, cat # 07183.

### PylRS library screen in E. coli

A plasmid library composed of the 20 canonical amino acids randomized at 5 sites was used alongside positive (pREP-pylT) and negative selection (pYOBB2-pylT) plasmids to select for aminoacyl-tRNA synthetases unique to pMe-2,3,5,6-tetrafluoro-Phe. All incubations for selection are carried out at 37ºC for 16 hours unless otherwise noted. All recovery steps last 1 hour and are shaken at 250 rpm. Cultures are grown at 37ºC for 16 hours and shaken at 250 rpm.

For positive selection, 500 µL of DH10B electrocompetent cells containing the positive pREP-pylT selection plasmid were transformed by electroporation with 1µg of library plasmid. Cells were recovered in SOC media at 37ºC for 1 hour. Recovered cells were serially plated from 10 to 10^−6^, and grown for 16 hours at 37ºC to insure proper library coverage. Coverage was calculated to ~150x, representing >99% of possible library members. The pooled recovery was used to inoculate 500 mL of LB media with 50 µg/mL of Kanamycin (Kan) and 25 µg/mL of tetracycline (Tet). The culture was grown overnight to saturation. This saturated culture was used to inoculate 500 mL of fresh LB media. This was grown to an OD_600_ of 4.1, and 100 µL was plated on each of ten 15 cm LB-agar plates containing 50 µg/mL Kan, 25 µg/mL Tet, 40 µg/mL chloramphenicol (Cm), and 1mM ncAA. Plates were incubated overnight. To harvest the resulting cells, 5 mL of LB-media was added to each plate. Colonies were then scraped from plates, pooled, and recovered for 1 hour. Cells were then pelleted and plasmid DNA extracted using a Macherey-Nagel miniprep kit. Resulting DNA was then plasmid separated by isolating the library plasmid on a 0.8% agarose gel via agarose gel electrophoresis and extracting using the Thermo Scientific GeneJET gel extraction kit.

For negative selection, 100 ng of plasmid DNA from the positive selection was transformed into 50 µL of DH10B cells containing PYOBB2-pylT using electroporation and recovered in 1 mL SOC media. Following recovery, 100 µL was plated on each of 3 LB-agar plates containing 50µg/mL Kan, 25µg/mL Cm, and 0.2% arabinose. After 16 hours, cells were scraped and DNA prepped as described above.

To identify functional synthetase variants, 100 ng of the pBK DNA from negative selection was transformed into 25 µL of electrocompetent DH10B cells containing the reporter plasmid, pALS-pylT. This plasmid contains the sfGFP reporter with a TAG site at residue 150. In the presence of the selected ncAA, the TAG site will be suppressed, and the resulting colonies should appear green. Transformed cells were recovered in 1 mL SOC media. After recovery, cells were diluted by a factor of 100, and 100 µL of this dilution was plated on three 15 cm auto-inducing plates [40]. All plates contain 50 µg/mL Kan and 25 µg/mL Tet; two contain 1 mM ncAA and the other is left without ncAA as a control. Plates were incubated overnight. After 16 hours, plates were kept at RT for 24 hours to further develop. After fully maturated, colonies that appeared green were individually selected and used to inoculate 500 µL of non-inducing media [65] containing 50 µg/mL Kan and 25 µg/mL Tet in a 96-well block. This block was incubated for 20 hours at 37ºC shaking at 300 rpm. After adequate growth, 20 µL of each well was used to inoculate two 96-well blocks with 500 µL of auto-inducing media [40, 65] with 50 µg/mL Kan and 25 µg/mL Tet. One block contained 1 mM ncAA while the other did not. After 24 hours of incubation (37ºC, 300 rpm), sfGFP fluorescence was measured. The 30 highest performing synthetases were sequenced, and 17 unique sequences were identified.

### UP50 measurements

Electrocompetent DH10B cells containing the pALS reporter plasmid were transformed with isolated pBK plasmids containing the Phe_X_ synthetases. The transformants of 8 synthetases, A11, A12, B5, B6, C4, C10, D6 and D7, were used to inoculate five 500 µL cultures each, consisting of auto-induction media with 50 µg/mL Kan and 25 µg/mL Tet containing no ncAA, 0.1 mM, 0.2 mM, 0.5 mM, or 1 mM ncAA. These were performed in duplicate, and fluorescence and OD_600_ were measured after 24 hours.

### Permissivity screen (E. coli)

Electrocompetent DH10B cells containing the pALS reporter plasmid were transformed with isolated pBK plasmids containing the synthetases. Cultures of 5 mL were inoculated with single colonies of each synthetase in non-inducing media [40] with 50 µg/mL kanamycin and 25 µg/mL tetracycline and grown for 16 hours. Noncanonical amino acid stocks were made at 100x their final concentration. 96-well blocks were prepared with 500 µL of auto-induction media containing antibiotics, and ncAAs with a final working concentration of 1 mM were added to their respective wells. Each well was then inoculated with a corresponding synthetase, yielding a final 96-well block where each synthetase was tested against each amino acid of interest. Cultures were grown in duplicate and fluorescence measurements were taken at 24 hours. Previous work has shown that fluorination of aromatic residues encoded at position N150 does not substantially affect intrinsic sfGFP fluorescence [64].

### Expression and purification of sfGFP (E. coli)

Previously existing E. coli containing sfGFP150TAG and a given RS in non-inducing media were used to inoculate 5mL cultures of non-inducing media containing 50 µg/mL Kan and 25 µg/mL Tet. Cultures were incubated for 20 hours at 37ºC, shaking at 250 rpm. After adequate growth, 500 µL of the 5 mL cultures were used to inoculate 50mL cultures of auto-inducing media containing 1 mM ncAA. Cultures were incubated for 24 hours (37ºC, 250 rpm), after which cultures were then spun down at 5000 rcf for 10 minutes and resuspended in 10 mL of a Tris buffer solution containing 100mM Tris, 0.5 M NaCl, and 5mM imidazole. Cells were lysed by micro-fluidization at 18,000 psi and centrifuged at 20,0000 rcf for 30 minutes. sfGFP proteins in cell lysate were bound to TALON Metal Affinity Resin (bed volume of 100 µL) at 4ºC for 1 hour. Lysate and bound resin were transferred to a gravity-flow column and the resin was washed with 30 bed volumes of buffer. Protein was eluted using Tris buffer solution previously described supplemented with 200 mM imidazole. Protein concentration was assessed by comparing sfGFP fluorescence to a standard curve and submitted for ESI-MS.

As is apparent in the traces shown in Figure 4, the deconvoluted mass spectra showed multiple masses for WT as well as mutant sfGFP. The spectra were analyzed in-house by Novatia (Promass) to yield relative intensities at given masses. In addition to the base (main) peak, both WT and mutant sfGFP displayed minor masses of ~-131 Da (lacking Methionine 1), ~-1378 Da, and ~-4165 Da. For the purpose of estimating encoding fidelity, these were considered part of the expected mass, and all other signals were considered non-expected mass. Note that the first time that sfGFP-N150TAG was expressed via Phe_X-D6_ + 2,3,6F Phe, a minor +36.9 Da peak was present in the spectra. This was classified as a K+ adduct of the expected mass by Promass software. It was suggested to us that this mass may have in fact been due to contamination of this sample by sfGFP with penta-fluoro Phe encoded (as a result of technical error). We re-expressed the sfGFP-N150TAG via Phe_X-D6_ + 2,3,6F Phe, and in this second purification we did not observe the +36.9 Da peak. The trace for this sample is shown in Figure 4. For Phe_X-D6_, encoding fidelity for Penta-, 2,3,5,6-, 2,3,6- and 2,6-flouro Phe was 98.2%, 98.7%, 100.0%, and 95.0% respectively. For Phe_X-B5_, encoding fidelity for Penta-, 2,3,5,6-, 2,3,6-, 2,6-, and 2-fluoro Phe was 97.5%, 97.9%, 88.1%, 81.8%, and 95.6%.

### Mammalian Cell culture

HEK 293T cells (CRL 3216) were maintained in DMEM high glucose medium supplemented with Pen/Strep, L glutamine, and 10% FBS (Sigma). Cells were used at passages 5 to 35.

### Construction of pAcBac1-Phe_X_ Plasmid

The pAcBac1.tR4-MbPyl, used to express a MbPylRS in mammalian cells, was a gift from Peter Schultz (Addgene plasmid # 50832). A WT human codon optimized version of the MbPylRS was substituted into the pAcBac1.tR4-MbPyl and was a gift from Jason Chin. The active site sequence with mutated residues was synthesized as a gBlock from Integrated DNA Technologies. Relevant DNA fragments were assembled using NEBuilder HiFi DNA Assembly Master Mix, transformed and isolated from NEB Stable cells, and sequence confirmed.

### Permissivity screen (mammalian)

HEKT cells were seeded in 60 mm dishes so that they would be approximately 60-80% confluent the next day. We transfected the cells using 1.75 µg of the PacBac1-based synthetase plasmid (Phe_X-D6_ or Phe_X-B5_) and 0.75 µg of sfGFP_N150TAG per dish. Polyjet was used (7.5 µl per dish, 2.5 mL media volume). Media was exchanged to contain ncAAs at a given concentration before transfection and approximately 16 hours after transfection. Amino acids were solubilized either in NaOH for those synthesized as free acids (penta-flouro, 2,3,6 trifluoro, 3,4,5 trifluoro, 2,5 difluoro, 2,4 difluoro, 2 mono) or in HCl for those synthesized as HCl salts (2,3,5,6 tetrafluoro, 2,3,4,5 tetrafluoro, 2,6 difluoro). Cells were imaged and harvested approximately 24 hrs after transfection. Images were taken with 20X objective using a Leica DFC9000GT camera with 700 ms exposure, without binning. Excitation was through an X-Cite 120 LED (Lumen Dynamics) set to 100% intensity. For bright field images, light was adjusted manually. To harvest, cells were washed twice with ice cold DPBS, then sloughed off the dishes in 1 mL ice cold DPBS supplemented with Roche protease inhibitors. Cells were collected in microcentrifuge tubes and pelleted by centrifugation. Supernatant was removed and the cell pellets were flash frozen in liquid nitrogen, then stored at -80C. To lyse, 350 ul of RIPA buffer (Sigma) plus Roche protease inhibitor tablets was added to each pellet. After cell debris and nuclei were cleared via centrifugation, GFP fluorescence was read from supernatants in 96 well plates; for each concentration of amino acid, at least 6 readings were made from at least 2 transfections. The level of fluorescence from lysates from untransfected cells averaged 21.7 ± 2.1 RFU. By comparison, cells transfected with sfGFP_N150TAG and Phe_X-D6_ averaged 26.9 ± 5.6 RFU and Phe_X-B5_ averaged 43.1 ± 2.5. The maximum background signal thus attributable to Phe_X-B5_ and Phe_X-D6_ was thus ~5 RFU and ~21.4 RFU respectively.

### Expression for purification and mass spectrometry (mammalian)

HEK 293T cells were expanded in 10 mm dishes. For a given synthetase/amino acid combination, the size of the transfection (number of plates) was estimated from differences in relative sfGFP_N150TAG yield in the multi-specificity screens, with poorer encoders requiring more cell mass. Overall, 7-40 dishes were transfected with master mixes such that each dish received 3.5 µg of pACBAC1-Phe_x_, 1.5 µg sfGFP_N150TAG, and 15 µL of PolyJET according to manufacturer’s instructions. Media was changed before transfection and approximately 18 hrs after transfection. Cells were harvested approximately 36 hours after transfection. To harvest, each dish was washed twice with DPBS. Cells were harvested via scraping in ice-cold DPBS plus Roche protease inhibitors, on ice, pelleted by centrifugation, and flash frozen in liquid nitrogen. Cells were lysed in hypotonic lysis buffer (10 mM Tris/HCl pH 8, Roche protease inhibitors, 0.1% Triton X 100 Sigma) via dounce homogenization on ice. Cell debris and nuclei were cleared via centrifugation. Clarified lysate was diluted 1:2 in Wash buffer (25 mM Tris/HCl pH 8, 20 mM Imidazole, 150 mM NaCl) and applied to pre-equilibrated Ni-NTA resin (Qiagen). Column was washed in at least 30 volumes of wash buffer, then protein was eluted in elution buffer (25 mM Tris/HCl pH 8, 250 mM Imidazole, 150 mM NaCl). Elution was exchanged into 100 mM TEAB, prepared via 1:10 dilution of 1 M Stock (thermofisher) in ultrapure water. Finally, protein was concentrated to ~0.5 mg/ml, flash frozen, and submitted to Novatia Inc for ESI mass spec of the intact protein. The LC-UV traces from Novatia indicated that purified WT and mutant sfGFP existed as three predominant isoforms, a dominant peak and two minor peaks representing +532 +/-1 Da and -42 +/-1 Da mass differences (Supplemental Fig 4). Note that the WT condition was co-transfected with Phe_X-B5_ synthetase, which did not affect the mass (within <1 Da of the same WT sfGFP expressed alone [40]). As was also the case when expressed in *E*.*coli*, the deconvoluted mass spectra for the dominant LC peak showed multiple masses for WT, as well as mutant sfGFP, from HEK cells. The spectra was analyzed in-house by Novatia (Promass software) to yield relative intensities of the dominant mass and any other apparent peaks. WT and mutant sfGFP shared minor peaks of ~ -42 Da (appx 2% of integration), ~ -1288 Da (appx 2% of integration) ~183 Da (appx 3% of integration), and ~ +53 Da (appx 1% of integration) in common and thus, for the sake of estimation of encoding fidelity, these were counted among the expected mass. To be conservative, all other peaks were considered unexpected mass, even though they may not represent actual misincorporation. Fidelity for Phe_X-D6_ and Penta-, 2,3,4,5-, 2,3,5,6-, 2,3,6-, 2,6-,and 2,5-fluoro phenylalanine were estimated to be 97.5%, 100%, 98.2%, 98.5%, 97.6%, and 95.6%. For Phe_X-B5_, fidelity was 96.2%, 97.4%, 100%, 100%, 100%, and 91.3% for the same species.

To accurately estimate sfGFP yield in a scenario wherein loss during purification is minimized, we transfected and harvested 31-10 cm dishes of HEKT cells with Phe_X-D6_ and sfGFP_N150TAG in the presence of 2 mM penta-fluoro Phenylalanine. sfGFP was purified as described above. Percent sfGFP was quantified using band detection and densitometry in Biorad Imagelab software (68% of lane signal).

### Expression of ion channel constructs with site-specific encoding of 2,3,6F Phe

pCMV-CFTR was a gift from Paul McCray (University of Iowa). WT CFTR-TAA (the existing TAG stop codon changed to TAA), F508 TAG CFTR (TAG amber codon introduced at F508 in the above WT construct), as well as WT and F1486TAG hNav 1.5_strep_HIS_strep were produced using standard methods and sequenced through the open reading frame. For expression of WT CFTR, 10 cm dishes of HEK 293T cells were transfected with 1.5 µg WT CFTR-TAA and 0.25 µg WT GFP. For expression of WT hNav 1.5, 10 cm dishes of HEK 293T cells were transfected with 1.5 µg of WT hNav 1.5_strep_HIS_strep and 0.25 µg WT GFP. For encoding of 2,3,6F Phe, 10 cm dishes of HEK 293T cells were transfected from master mixes of DNA such that each dish received 3.5 µg of Phe_X-D6_ and 0.25 µg of sfSFGFP_N150TAG. Depending on condition, 2 µg of F508TAG-CFTR or 1.5 µg of F1486TAG-Nav 1.5-SHS were added. Media was changed twice (~16 hr and ~32 hr post transfection), and cells were harvested ~44 hrs post transfection and flash frozen in aliquots. Cell pellets were resuspended in RIPA (Sigma Aldrich) plus Roche protease inhibitors. 60 µg of protein from cleared, unconcentrated lysate (or a dilution as indicated) was loaded on 4-20% SDS gels. For Western blotting, gels were transferred to Nitrocelllose membranes and probed with anti-CFTR (ab596 from Cystic Fibrosis Foundation, 1:1750) or anti-Nav 1.5 (D9J7S, Cell Signaling, 1:5000) overnight in 5% milk in TBST. Blots were washed, probed with HRP secondaries (1:10,000) and imaged using Clarity ECL (Biorad). Blots were subsequently probed for beta-actin using HRP-conjugated primary (AC-15, Novus Biologicals 1:2000) appx 3 hours in 5% milk in TBST. Blots were washed and imaged using Clarity ECL (Biorad). Densitometry (background subtracted integrated density) was done in ImageJ.

For patch clamp recording of hNa_v_ 1.5, in parallel we co-transfected 10 cm dishes with 3.5 µg of Phe_X-D6_, 0.25 µg of sfGFP_N150TAG, and 1.5 µg of either WT hNa_v_ 1.5 strep_HIS_strep or hNa_v_ 1.5 F1486TAG strep_HIS_strep. Both conditions were cultured in 2 mM 2,3,6 Trifluoro Phe throughout. Media was changed at 16 hrs and again when seeding onto 35 mm dishes for recording. Recordings were done approximately ~42 hrs post transfection. Pipette solution contained 105 CsF, 35 NaCl, 10 EGTA, and 10 Hepes (pH 7.4). The bath contained (in mmol/L) 150 NaCl, 2 KCl, 1.5 CaCl2, 1 MgCl2, and 10 Hepes (pH 7.4). Pipette resistances were around 2 mΩ and series resistance was compensated ≥ 85%. For generation of I/V curves, cells were held at -140 mV and pulsed in 5 mV depolarizing increments. For measurement of steady state inactivation, cells were held at -140 mV and conditioned for 500 ms in 5 mV depolarizing increments. Test pulse was at -30 mV. Normalized G/V and SSI curves were fit to Boltzmann functions using Origin software.

For purification of hNa_v_ 1.5 F1486(2,3,6F Phe), ten 10 cm dishes of HEKT cells were co transfected so that each dish received 1.5 µg pCDNA hNa_v_ 1.5 F1486TAG_FLAG, 3.5 µg Phe_X-D6_, and 0.25 µg sfGFPN150TAG plasmids. Media changes were as described for the western blots. Channels were solubilized and purified as described in the report of the cryo-EM structure of the cardiac channel [47], with the following modifications. First, we used DDM/CHS as detergent throughout rather than exchanging to GDN during washes. Secondly, the elution was concentrated and run directly on SDS page gel instead of being subjected to size exclusion chromatography. Gel band excision, tryptic digestion, and MS/MS were performed by the Iowa Proteomics Core using standard methods.

### Quantum calculations of cation pi binding potential

All the models are prepared using GaussView 6 and all quantum chemical calculations are performed using Gaussian 16 [66]. Full geometry optimizations are carried out at M06/6-31G(d,p) level of theory. In some fluorinated aromatic cases where the optimization resulted in geometries that are not considered cation-*π* interactions, the binding energy is obtained using the single-point method, as done previously [25]. In this approach full geometry optimization of the cation and the aromatic system is performed without the fluorination at the position which caused incompatible geometries upon optimization. Then fluorine is appended to the aromatic ring with a bond distance calculated from the aromatic system with fluorination at the desired position which is optimized in isolation. The energy of this system is calculated and used to find the binding energy in the single-point calculations.

## Supplemental Figures

**Supplemental Fig 1:**
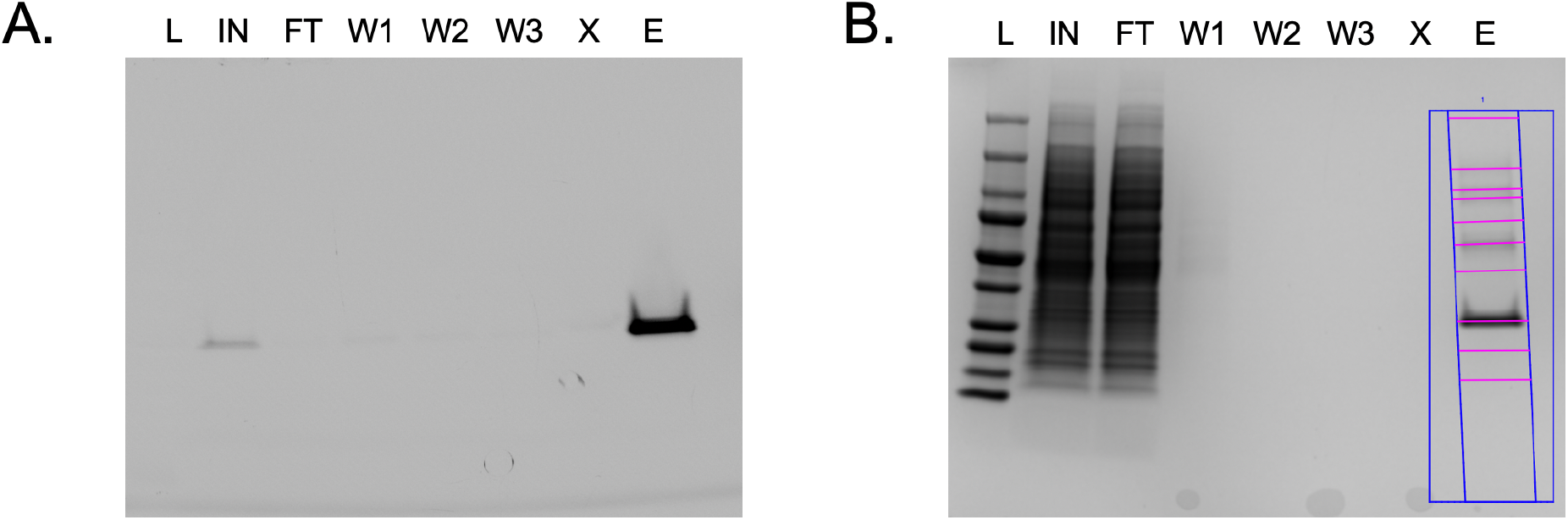
Large scale prep and yield of sfGFP bearing penta-F phenylalanine at position N150 using Phe_x.D6_-Purification was started from 3.26 g of HEK cells (31 10 cm dishes) **A**. SDS page gel imaged for GFP before staining L= Ladder, IN= input, i.e., diluted lysate, FT= flowthrough, W1, W2, W3 = washes, X= empty, E = elution (diluted 1:1 in wash buffer). 30 **µ**L of sample loaded per lane from 100 mL (IN/FT), 10 mL (Washes), 1.5 mL (E). Thus, ∼1/50 of the elution was run in lane E. **B**. Coomassie stain of the same gel. Buffer exchange and concentration of elution resulted in 240 **µ**L of 1.041 abs. Factoring the extinction coefficient of sfGFP_V5_HIS and a purity estimate (68%) from densitometry of the bands in Coomassie of E yields a total sfGFP estimate of 110.4 **µ**g, or 34 **µ**g/g of cells (wet pellet weight).

**Supplemental Figure 2.**
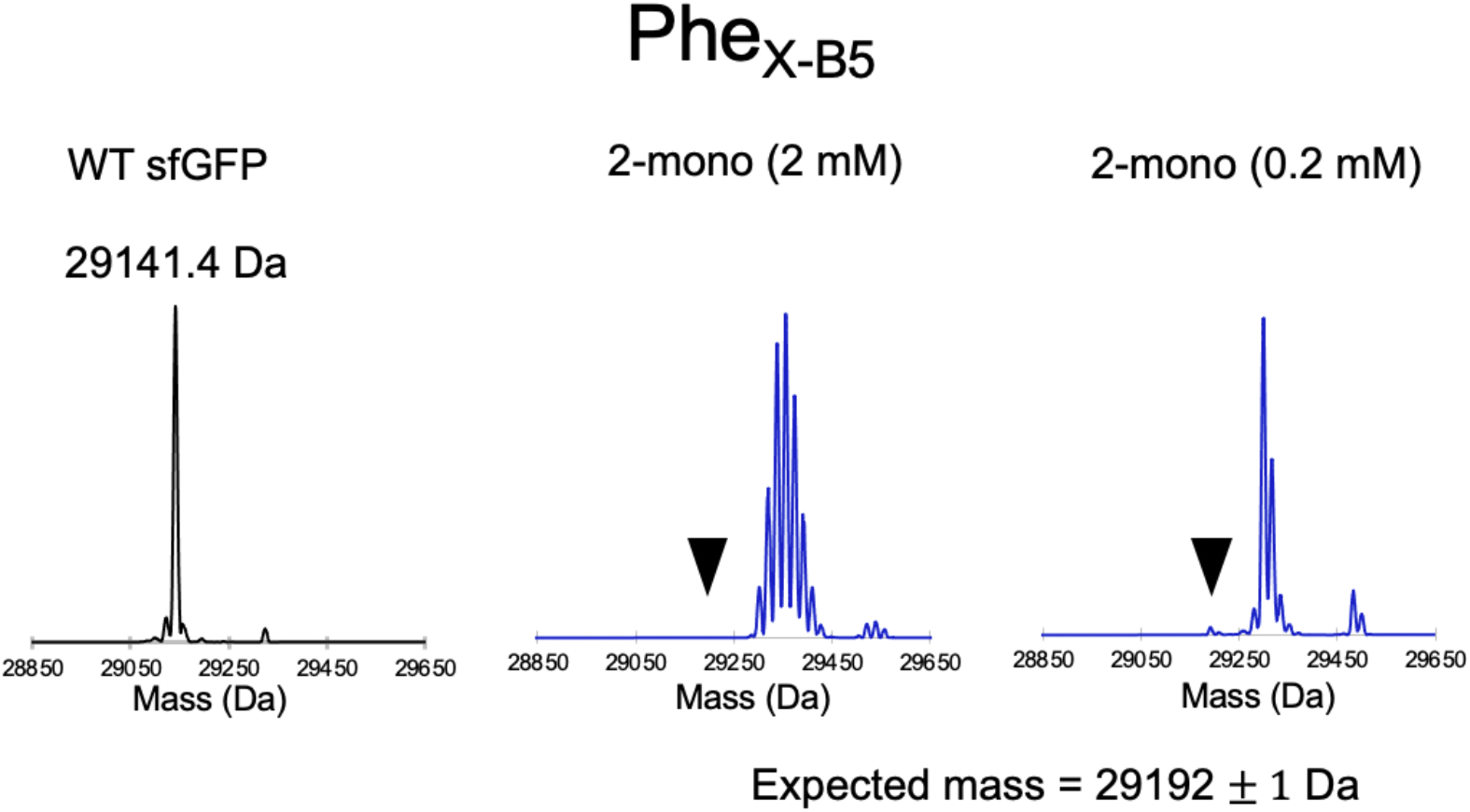
Deconvoluted ESI mass spectra for sfGFP-N150TAG expressed with 2-mono fluoro phenylalanine and PheX-B5. Both 2 mM and 0.2 mM concentration of amino acid resulted in extensive incorporation of the amino acid at natural Phe codons, as indicated by masses at multiples of +18 Da. Signal at expected mass is absent in the 2 mM condition and a very small fraction (∼1%) in 0.2 mM condition (black arrowheads).

**Supplemental Figure 3:**
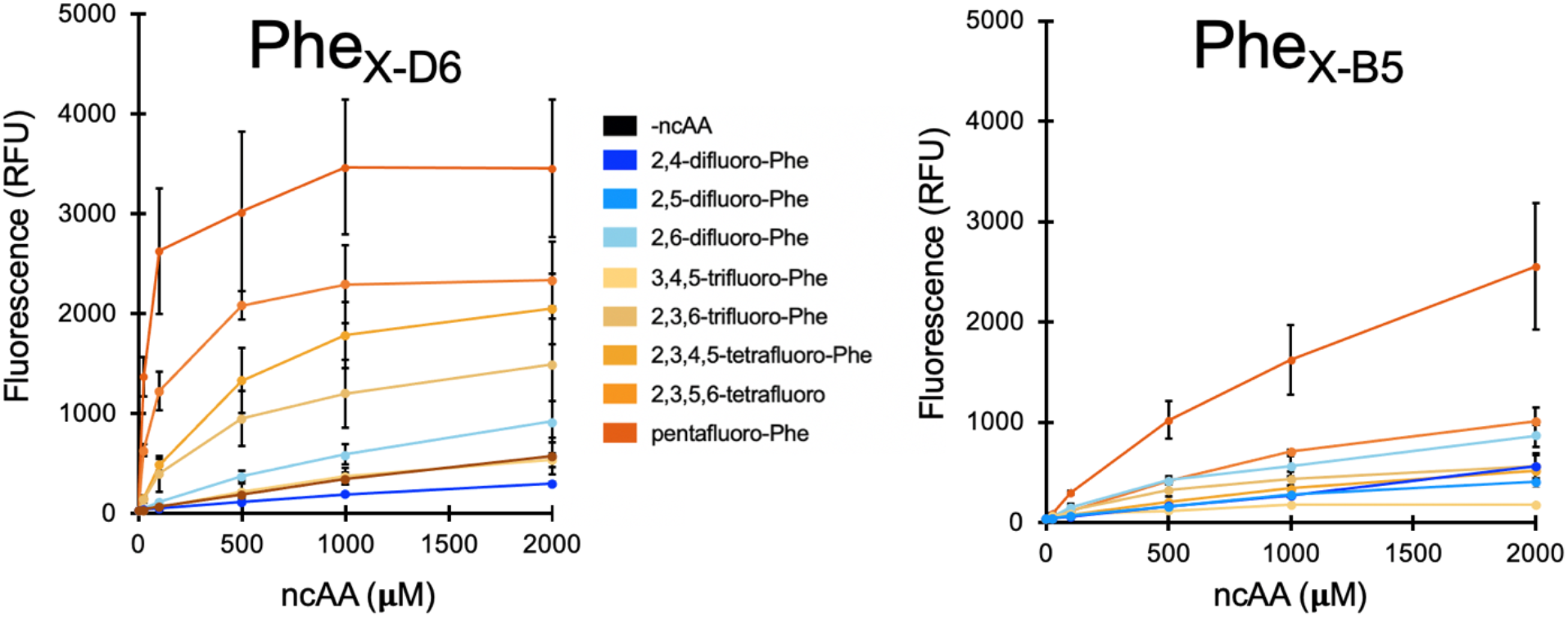
GFP lysate values from HEKT cells co transfected with sfGFP150TAG and either Phe_x.D6_ (left) or Phe_x.B5_ (right) and harvested approximately 24 hours post transfection. Error bars are ± STD.

**Supplemental Figure 4:**
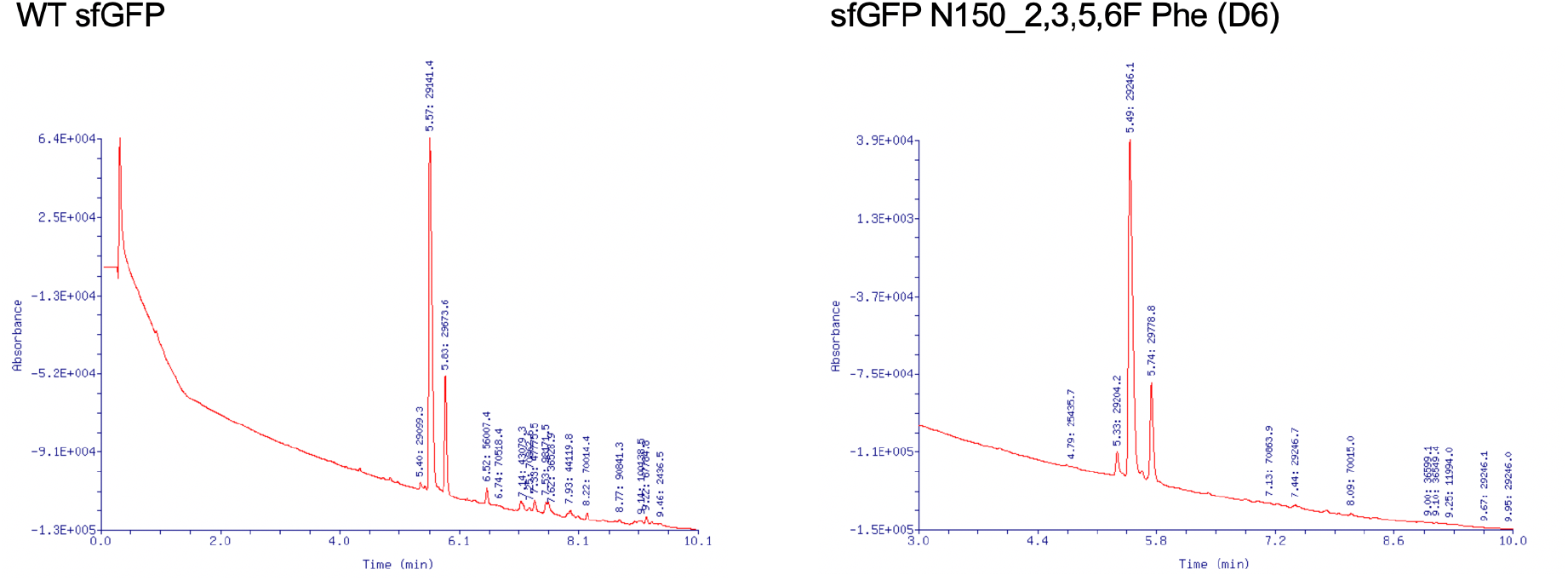
UV absorbance of Liquid Chromatography of purified sfGFP samples. Both WT (left) and mutants (in this case D6 encoding 2,3,5,6 tetra-F Phe) eluted as 3 typical peaks for sfGFP, a dominant peak, as well as one at ∼ -42 Da and a third at ∼ + 532 Da. The deconvoluted mass spectra for the dominant (center) peaks are shown in the main figures.

**Supplemental Figure 5:**
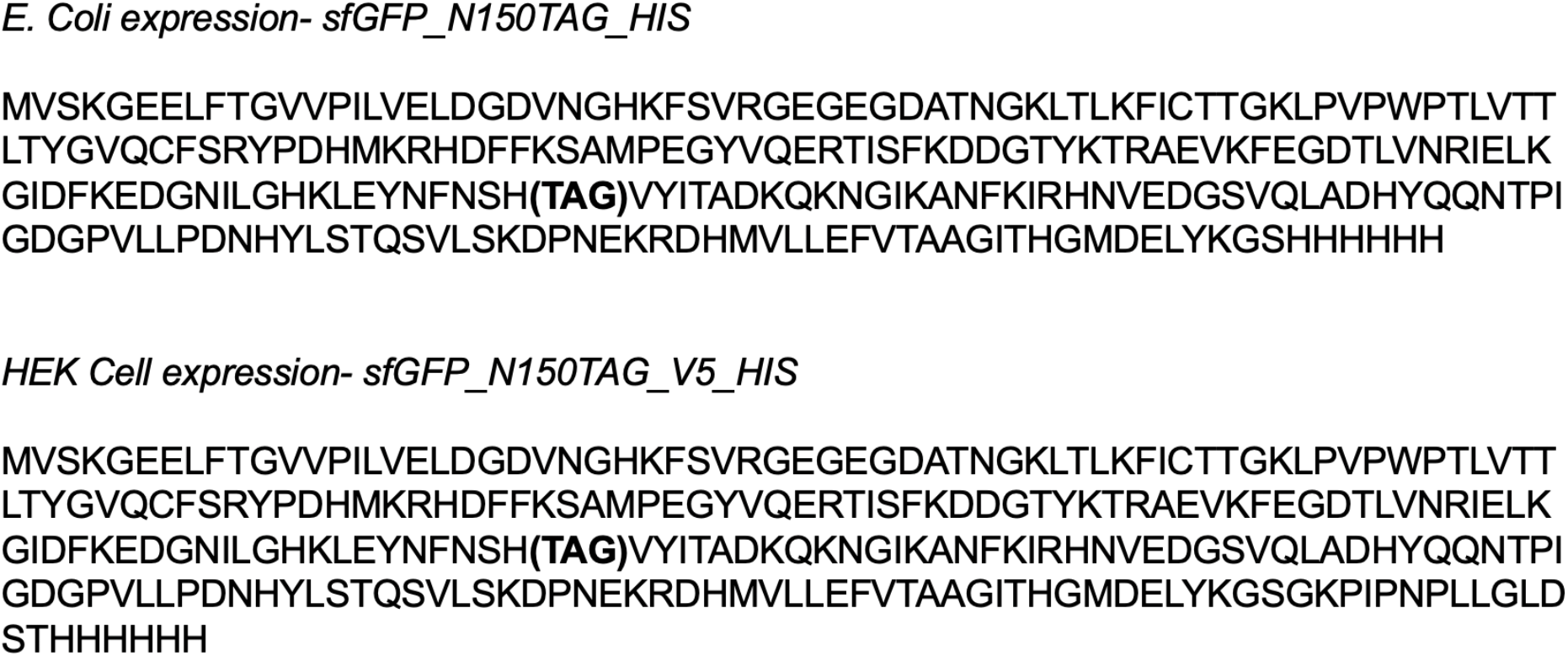
Protein sequences of sfGFP variants used for expression and purification in this study.

**Supplemental Table 1:** Gating Parameters for hNa_v_ 1.5 variants. **Bolded** = p value of 0.0014 compared to WT. P>0.24 for all other comparisons.

|  | Activation |  |  |  |  | Steady State Inactivation |  |  |  |  |
| --- | --- | --- | --- | --- | --- | --- | --- | --- | --- | --- |
|  | N | V<sub>mid</sub> | SEM | d(x) | SEM | N | V<sub>mid</sub> | SEM | d(x) | SEM |
| WT<br>hNav1.5 | 5 | -52.0 | 1.71 | 7.37 | 0.58 | 5 | -97.8 | 0.85 | 6.76 | 0.45 |
| F1486<br>(2,3,6F) | 8 | -52.8 | 1.56 | 7.31 | 0.66 | 8 | **-104.5** | 1.33 | 7.43 | 0.25 |

## Acknowledgements

GDG, DTI, CJC, SM, JDG and CAA were supported by NIGMS R01106569 and NINDS R24 NS104617 RBC and RTM were supported in part by the GCE4All Biomedical Technology Development and Dissemination Center supported by National Institute of General Medical Science grant RM1-GM144227 ET and AR were supported by NIH P41-104601

## References

1. Dougherty, D.A., Cation-pi interactions in chemistry and biology: a new view of benzene, Phe, Tyr, and Trp. Science, 1996. 271(5246): p. 163–8.

2. Schottel, B.L., H.T. Chifotides, and K.R. Dunbar, Anion-pi interactions. Chem Soc Rev, 2008. 37(1): p. 68–83.

3. Dougherty, D.A., The cation-pi interaction. Acc Chem Res, 2013. 46(4): p. 885–93.

4. Weber, D.S. and J.J. Warren, The interaction between methionine and two aromatic amino acids is an abundant and multifunctional motif in proteins. Arch Biochem Biophys, 2019. 672: p. 108053.

5. Ringer, A.L., A. Senenko, and C.D. Sherrill, Models of S/pi interactions in protein structures: comparison of the H2S benzene complex with PDB data. Protein Sci, 2007. 16(10): p. 2216–23.

6. Daeffler, K.N., H.A. Lester, and D.A. Dougherty, Functionally important aromatic-aromatic and sulfur-pi interactions in the D2 dopamine receptor. J Am Chem Soc, 2012. 134(36): p. 14890–6.

7. Burley, S.K. and G.A. Petsko, Amino-aromatic interactions in proteins. FEBS Lett, 1986. 203(2): p. 139–43.

8. Liao, S.M., et al., The multiple roles of histidine in protein interactions. Chem Cent J, 2013. 7(1): p. 44.

9. Levitt, M. and M.F. Perutz, Aromatic rings act as hydrogen bond acceptors. J Mol Biol, 1988. 201(4): p. 751–4.

10. Pace, C.J. and J. Gao, Exploring and exploiting polar-pi interactions with fluorinated aromatic amino acids. Acc Chem Res, 2013. 46(4): p. 907–15.

11. Lucas, X., et al., A thorough anion-pi interaction study in biomolecules: on the importance of cooperativity effects. Chem Sci, 2016. 7(2): p. 1038–1050.

12. Infield, D.T., et al., Cation-pi Interactions and their Functional Roles in Membrane Proteins. J Mol Biol, 2021. 433(17): p. 167035.

13. Gallivan, J.P. and D.A. Dougherty, Cation-pi interactions in structural biology. Proc Natl Acad Sci U S A, 1999. 96(17): p. 9459–64.

14. Yen, T.J., et al., Structure of the saxiphilin:saxitoxin (STX) complex reveals a convergent molecular recognition strategy for paralytic toxins. Sci Adv, 2019. 5(6): p. eaax2650.

15. Rahman, M.M., et al., Structure of the Native Muscle-type Nicotinic Receptor and Inhibition by Snake Venom Toxins. Neuron, 2020. 106(6): p. 952–962 e5.

16. Plenge, P., et al., The antidepressant drug vilazodone is an allosteric inhibitor of the serotonin transporter. Nat Commun, 2021. 12(1): p. 5063.

17. Silberberg, J.M., et al., Deciphering ion transport and ATPase coupling in the intersubunit tunnel of KdpFABC. Nat Commun, 2021. 12(1): p. 5098.

18. Yang, D. and E. Gouaux, Illumination of serotonin transporter mechanism and role of the allosteric site. Sci Adv, 2021. 7(49): p. eabl3857.

19. Kumar, A., et al., Mechanisms of activation and desensitization of full-length glycine receptor in lipid nanodiscs. Nat Commun, 2020. 11(1): p. 3752.

20. McDaniel, M.J., et al., NMDA receptor channel gating control by the pre-M1 helix. J Gen Physiol, 2020. 152(4).

21. Kise, Y., et al., Structural basis of gating modulation of Kv4 channel complexes. Nature, 2021. 599(7883): p. 158–164.

22. Diwanji, D., et al., Structures of the HER2-HER3-NRG1beta complex reveal a dynamic dimer interface. Nature, 2021. 600(7888): p. 339–343.

23. Waheed, Q., et al., Interfacial Aromatics Mediating Cation-pi Interactions with Choline-Containing Lipids Can Contribute as Much to Peripheral Protein Affinity for Membranes as Aromatics Inserted below the Phosphates. J Phys Chem Lett, 2019. 10(14): p. 3972–3977.

24. Last, N.B., et al., Molecular determinants of permeation in a fluoride-specific ion channel. Elife, 2017. 6.

25. Davis, M.R. and D.A. Dougherty, Cation-pi interactions: computational analyses of the aromatic box motif and the fluorination strategy for experimental evaluation. Phys Chem Chem Phys, 2015. 17(43): p. 29262–70.

26. Mecozzi, S., A.P. West, and D.A. Dougherty, Cation-pi interactions in simple aromatics: Electrostatics provide a predictive tool. Journal of the American Chemical Society, 1996. 118(9): p. 2307–2308.

27. Leisle, L., et al., Incorporation of Non-Canonical Amino Acids. Adv Exp Med Biol, 2015. 869: p. 119–51.

28. Santarelli, V.P., et al., A cation-pi interaction discriminates among sodium channels that are either sensitive or resistant to tetrodotoxin block. J Biol Chem, 2007. 282(11): p. 8044–51.

29. Santarelli, V.P., et al., Calcium block of single sodium channels: role of a pore-lining aromatic residue. Biophys J, 2007. 93(7): p. 2341–9.

30. Pless, S.A., et al., Molecular basis for class Ib anti-arrhythmic inhibition of cardiac sodium channels. Nat Commun, 2011. 2: p. 351.

31. Ahern, C.A., et al., Electrostatic contributions of aromatic residues in the local anesthetic receptor of voltage-gated sodium channels. Circ Res, 2008. 102(1): p. 86–94.

32. Zhong, W., et al., From ab initio quantum mechanics to molecular neurobiology: a cation-pi binding site in the nicotinic receptor. Proc Natl Acad Sci U S A, 1998. 95(21): p. 12088–93.

33. Padgett, C.L., et al., Unnatural amino acid mutagenesis of the GABA(A) receptor binding site residues reveals a novel cation-pi interaction between GABA and beta 2Tyr97. J Neurosci, 2007. 27(4): p. 886–92.

34. Budisa, N., Engineering the genetic code : expanding the amino acid repertoire for the design of novel proteins. 2006, Weinheim: Wiley-VCH. xvi, 296 p.

35. Brown, W., J. Liu, and A. Deiters, Genetic Code Expansion in Animals. ACS Chem Biol, 2018. 13(9): p. 2375–2386.

36. Lee, Y.J., et al., Genetically encoded fluorophenylalanines enable insights into the recognition of lysine trimethylation by an epigenetic reader. Chem Commun (Camb), 2016. 52(85): p. 12606–12609.

37. Hao, B., et al., A new UAG-encoded residue in the structure of a methanogen methyltransferase. Science, 2002. 296(5572): p. 1462–6.

38. Arbely, E., et al., Photocontrol of tyrosine phosphorylation in mammalian cells via genetic encoding of photocaged tyrosine. J Am Chem Soc, 2012. 134(29): p. 11912–5.

39. Srinivasan, G., C.M. James, and J.A. Krzycki, Pyrrolysine encoded by UAG in Archaea: charging of a UAG-decoding specialized tRNA. Science, 2002. 296(5572): p. 1459–62.

40. Galles, G.D., et al., Selection and validation of orthogonal tRNA/synthetase pairs for the encoding of unnatural amino acids across kingdoms. Methods Enzymol, 2021. 654: p. 3–18.

41. Cooley, R.B., P.A. Karplus, and R.A. Mehl, Gleaning unexpected fruits from hard-won synthetases: probing principles of permissivity in non-canonical amino acid-tRNA synthetases. Chembiochem, 2014. 15(12): p. 1810–9.

42. Zheng, H., K. Comeforo, and J. Gao, Expanding the fluorous arsenal: tetrafluorinated phenylalanines for protein design. J Am Chem Soc, 2009. 131(1): p. 18–9.

43. Chatterjee, A., et al., Efficient viral delivery system for unnatural amino acid mutagenesis in mammalian cells. Proc Natl Acad Sci U S A, 2013. 110(29): p. 11803–8.

44. Rowe, S.M., S. Miller, and E.J. Sorscher, Cystic fibrosis. N Engl J Med, 2005. 352(19): p. 1992–2001.

45. Rivaud, M.R., M. Delmar, and C.A. Remme, Heritable arrhythmia syndromes associated with abnormal cardiac sodium channel function: ionic and non-ionic mechanisms. Cardiovasc Res, 2020. 116(9): p. 1557–1570.

46. Remme, C.A. and C.R. Bezzina, Sodium channel (dys)function and cardiac arrhythmias. Cardiovasc Ther, 2010. 28(5): p. 287–94.

47. Jiang, D., et al., Structure of the Cardiac Sodium Channel. Cell, 2020. 180(1): p. 122–134 e10.

48. Cai, Z., et al., Application of high-resolution single-channel recording to functional studies of cystic fibrosis mutants. Methods Mol Biol, 2011. 741: p. 419–41.

49. Gunderson, K.L. and R.R. Kopito, Conformational states of CFTR associated with channel gating: the role ATP binding and hydrolysis. Cell, 1995. 82(2): p. 231–9.

50. Denning, G.M., et al., Processing of mutant cystic fibrosis transmembrane conductance regulator is temperature-sensitive. Nature, 1992. 358(6389): p. 761–4.

51. Chakrabarti, S., et al., MOG1 rescues defective trafficking of Na(v)1.5 mutations in Brugada syndrome and sick sinus syndrome. Circ Arrhythm Electrophysiol, 2013. 6(2): p. 392–401.

52. Liu, F., et al., Structural identification of a hotspot on CFTR for potentiation. Science, 2019. 364(6446): p. 1184–1188.

53. Pan, X., et al., Structure of the human voltage-gated sodium channel Nav1.4 in complex with beta1. Science, 2018. 362(6412).

54. Shen, H., et al., Structural basis for the modulation of voltage-gated sodium channels by animal toxins. Science, 2018. 362(6412).

55. Liu, F., et al., Molecular Structure of the Human CFTR Ion Channel. Cell, 2017. 169(1): p. 85–95 e8.

56. Zhang, Z. and J. Chen, Atomic Structure of the Cystic Fibrosis Transmembrane Conductance Regulator. Cell, 2016. 167(6): p. 1586–1597 e9.

57. Laselva, O., et al., Identification of binding sites for ivacaftor on the cystic fibrosis transmembrane conductance regulator. iScience, 2021. 24(6): p. 102542.

58. Veit, G., et al., Allosteric folding correction of F508del and rare CFTR mutants by elexacaftor-tezacaftor-ivacaftor (Trikafta) combination. JCI Insight, 2020. 5(18).

59. Serohijos, A.W., et al., Phenylalanine-508 mediates a cytoplasmic-membrane domain contact in the CFTR 3D structure crucial to assembly and channel function. Proc Natl Acad Sci U S A, 2008. 105(9): p. 3256–61.

60. Fiedorczuk, K. and J. Chen, Mechanism of CFTR correction by type I folding correctors. Cell, 2022. 185(1): p. 158–168 e11.

61. Kellenberger, S., et al., Molecular analysis of the putative inactivation particle in the inactivation gate of brain type IIA Na+ channels. J Gen Physiol, 1997. 109(5): p. 589–605.

62. Danielson, M.A. and J.J. Falke, Use of 19F NMR to probe protein structure and conformational changes. Annu Rev Biophys Biomol Struct, 1996. 25: p. 163–95.

63. Awad, L.F. and M.S. Ayoup, Fluorinated phenylalanines: synthesis and pharmaceutical applications. Beilstein J Org Chem, 2020. 16: p. 1022–1050.

64. Miyake-Stoner, S.J., et al., Generating permissive site-specific unnatural aminoacyl-tRNA synthetases. Biochemistry, 2010. 49(8): p. 1667–77.

65. Studier, F.W., Protein production by auto-induction in high density shaking cultures. Protein Expr Purif, 2005. 41(1): p. 207–34002E

66. Frisch, M.J., et al., Gaussian 16 Rev. C.01. 2016: Wallingford, CT.

